# Prolonged exposure does not increase soil microbial community response to warming along geothermal gradients

**DOI:** 10.1101/102459

**Authors:** Dajana Radujković, Erik Verbruggen, Bjarni D. Sigurdsson, Niki I. W. Leblans, Sara Vicca, James T. Weedon

## Abstract

Predicting effects of climate change on ecosystem functioning requires knowledge of soil microbial community responses to warming. We used natural geothermal gradients (from +1°C to +19°C above ambient) in two subarctic grasslands to test the hypothesis that long-term exposure (>50 years) intensifies microbial community responses to warming compared to short-term exposure (5-7 years). Community profiles from amplicon sequencing of bacterial and fungal rRNA genes did not support this hypothesis: significant changes relative to ambient were observed from +9°C and upwards in the long-term and from 7°C to 11°C / +3°C to +5°C and upwards in the short-term, for bacteria and fungi, respectively. Our results suggest that bacterial communities in high-latitude grasslands will not undergo lasting shifts in community composition under the warming predicted for the coming 100 years. Fungal communities do appear to be temperature sensitive to the warming within this range, but only for short-term exposures.

## 1. INTRODUCTION

Given the recent concern regarding the stability of ecological systems under a changing climate, there have been numerous studies investigating the effects of warming on microbial community composition (Zogg et al. 1997; Zhang et al. 2005; Frey et al. 2008; Weedon et al. 2012; Luo et al. 2014; Karhu et al. 2014; Xu et al. 2015; Rui et al. 2015). The rationale behind these studies is that sensitivity of soil microbial community composition to increasing soil temperatures will indicate a corresponding sensitivity of the biogeochemical processes these communities are involved in (Zogg et al. 1997; Strickland et al. 2009).

Despite the large number of studies, a general conclusion as to how soil microbial communities will be affected by future temperature increases remains elusive. In some studies, short-term warming of approximately 1-2 years induced significant shifts in soil microbial community composition (Xiong et al. 2014; Zhang et al. 2016), while others did not find a significant change after substantially longer (4-9 years) warming treatments (Allison et al. 2010; Schindlbacher et al. 2011; Weedon et al. 2012). Rinnan et al. (2007) concluded that more than a decade of warming was needed to detect significant microbial responses in a manipulation experiment in sub-arctic heathlands. Similarly, DeAngelis et al. (2015), reported that microbial communities in a temperate forest were affected by a +5°C temperature increase only after 20 years of experimental warming. The lag in microbial response may occur because the substrate utilized by microbes shows a delayed response to warming and microbial communities will change only when substrate quantity and/or quality are significantly altered (Rinnan et al. 2007). Consistent with this, a meta-analysis by Blankinship et al. (2011) found that, for soil biota in general, the effects of warming intensified with time. Moreover, recent findings suggest that interannual variation in community composition might conceal warming effects in the short-term, but the apparent resistance may decline over time as warming continues to modify the soil substrate (Contosta et al. 2015). Therefore, it has been proposed that long-term warming experiments (on the scale of decades, rather than years) might be needed to detect the consequent changes in microbial community composition (Rinnan et al. 2007; Contosta et al. 2015). It can be assumed that microbial responses to warming over time would also depend on the warming magnitude given that any changes in the soil environment and microbial community composition should occur faster with increasing warming intensity.

Few studies have directly compared the effects of short-term and long-term warming of different intensities on soil microbial community composition. Such a comparison is crucial to assess whether responses observed in short-term experiments can be readily extrapolated to longer time scales (Rustad 2001; De Boeck et al. 2015). The present study tests the hypothesis that long-term warming (of several decades) intensifies microbial community response to warming so that a detectable change in community composition occurs at lower temperature elevations compared to short-term exposure (of several years) (Fig. 1). If this is the case then results observed over short timescales would potentially underestimate microbial response in the long-term.

**Figure 1.**
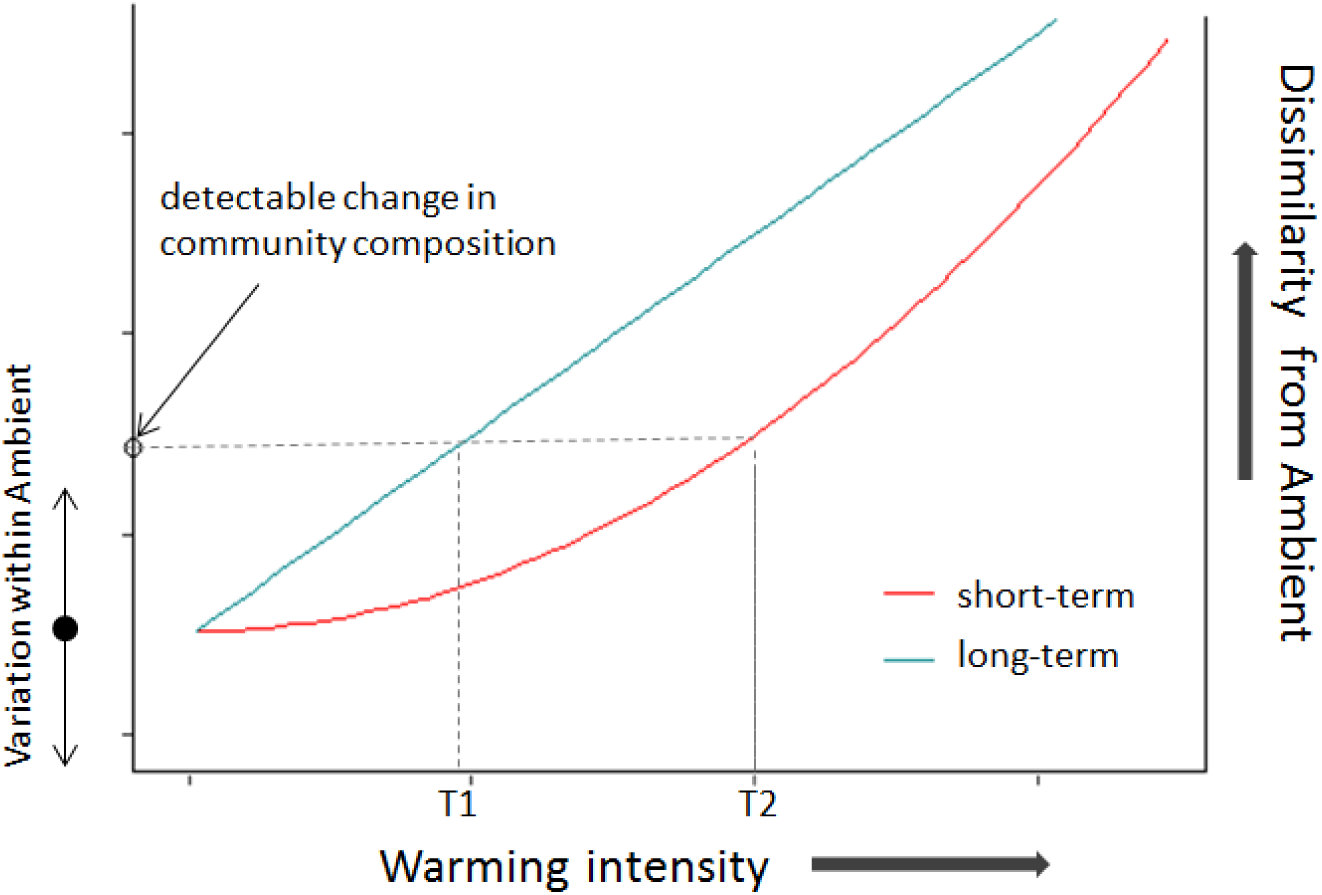
Microbial communities exposed to increasing temperature elevations are expected to become increasingly different from the ambient communities. However, natural and sampling variation, as well as a possible delay in the response of soil environment (substrate, vegetation) are expected to dampen this effect in the short-term and detectable effects of warming (where the community composition change surpasses the variation within ambient communities) occurs at higher temperatures (T2) than after a long period of warming where the soil environment is already significantly altered (T1). At very high temperature elevations, the curves for short- and long-term response are expected to converge, given that severe warming should produce fast responses. This model assumes the linear initial increase in dissimilarity between communities at increasing temperatures, however, similar explanation is applicable to other types of response curves (e.g. exponential, sigmoidal).

To test this hypothesis we took advantage of a “natural laboratory” afforded by a geothermal system in south-western Iceland. This area contains gradients in soil temperature of varying age (Sigurdsson et al. 2016), providing a unique opportunity to simultaneously investigate short- and long-term effects of warming within the same ecosystem type. Geothermal gradients have the advantage of providing a wide range of soil temperatures while avoiding the problems of both small-scale manipulative experiments, which are limited to a small number of temperature treatments; and large-scale latitudinal/altitudinal studies which often invoke difficulties in disentangling the effects of biogeography and temperature (Woodward et al. 2010; O’Gorman et al. 2014; De Boeck et al. 2015). Given that the geothermal gradients are contained within a relatively small area, potentially confounding environmental factors stay constant, allowing the effects of temperature to be isolated while at the same time retaining the complexity of the natural system (Woodward et al. 2010; O’Gorman et al. 2014).

## 2. MATERIALS AND METHODS

### 2.1 Site description and soil sampling

The study was conducted at ForHot research sites, which are located in the valleys surrounding the town Hveragerði (64°00′01″N, 21°11′09″W, 83-168 m a.s.l.) in south-western Iceland. Both sites represent unmanaged treeless grasslands dominated by the grass *Agrostis capillaris* L. The two grasslands contain two different geothermal systems; one has been geothermally heated for more than 50 years, (hereafter referred to as ‘long-term warmed grassland’ or LWG), the other has been exposed to warming since an earthquake that occurred on the 29^th^ of May 2008 (hereafter referred to as ‘short-term warmed grassland’ or SWG). After the earthquake in 2008, the locations of the SWG and LWG systems were mapped and it has been shown that, although geothermal manifestations in the LWG area were partly altered, the geothermal activity in the part of the LWG region where our study took place was not affected (Þorbjörnsson et al. 2009). Geothermal activity in the general area around LWG has most likely been persistent for several centuries, but the presence of multiple geothermal clay layers in few soil profiles at our LWG sites indicate that warming may have fluctuated somewhat in the past centuries. At both sites, the geothermal systems have resulted in gradients of soil temperature ranging from ambient (mean annual soil temperature of ≈5.1°C), to approximately +20°C (at a soil depth of 10 cm) over a distance of 50-100 m (see Sigurdsson et al. (2016) for more details on the temperature gradients).

The main soil characteristics of the two grasslands are comparable: soil type of both is Silandic Andosol; soil texture is silt loam; soil pH – 5.6 and 5.7; bulk density – 70 g/cm^3^ and 55 g/cm^3^ in SWG and LWG, respectively. pH, as an important factor influencing microbial community composition (Männistö et al. 2007), changes only slightly with increasing soil warming (from 5.6 to 6.3) (Sigurdsson et al. 2016).

In both grasslands, 5 replicate transects were established in autumn 2012. Each transect consists of 6 permanent plots (2m x 2m) located to span a range of different warming intensities: A≈+0°C (ambient), B≈+1°C, C≈+3°C, D≈+5°C, E≈+10°C and F≈+20°C. This system of plots was created as an attempt to make a comparable replicated warming gradient with matching temperature elevations in the two grasslands. However, because only instantaneous temperature was known when the plots were established and due to on-going small scale (spatial and temporal) fluctuations, the observed mean annual temperature deviated somewhat between matched plots across transects over longer time periods. Therefore we choose to re-classify the replicated plots based on their realized temperature differences. The range of actual warming in each of these new groups (averaged over the period May 2013 to May 2015) was as follows: W_0_ = 0 to +1°C (n = 11 and 10), W_low_ = +2 to +3°C (n = 5 and 4), W_med_ = +3 to +5°C (n = 4 and 6), W_high_ = +6 to +9°C (in LWG; n = 5) and W_high_’ = +7 to +11°C (in SWG; n = 5), W_extr_ = +15 to +19°C (n = 5 and 5 for LWG and SWG, respectively). There is thus, a slight difference between the grasslands in that LWG is slightly less warmed than SWG for W_high_ plots, whereas for other groups the two grasslands span the same range of warming intensities.

Soil samples were collected from all 60 plots in May 2013 for fungi and in July 2015 for bacteria, from 5-10 cm soil depth, using a corer (2.5 cm diameter). The fungal samples were collected and analysed in the framework of a study examining warming effects on SOM dynamics, the bacterial samples were collected as baseline measurements for an on-going multiyear fertilization experiment. Sampling protocols were identical on both sampling occasions. Individual soil samples were homogenized and stored at -20°C prior to further analyses.

### 2.2 Library preparation and sequencing

Total community DNA was extracted from approx. 0.25 g of soil using the PowerSoil DNA Isolation Kit according to the manufacturer’s protocol (MoBio, Carlsbad, CA, USA). For bacterial analyses, the hypervariable V3 region of the 16S rRNA gene was amplified using modified 341F and 518R primers (Bartram et al. 2011) with unique 6bp indices on the reverse primers. Each reaction mixture contained 1.5 μl of the sample, 1 μl each of forward and reverse primers (10μM) and 12.5 μl of Phusion High-Fidelity PCR Master Mix with HF Buffer (ThermoFisher Scientific, Waltham, MA, USA). PCR conditions were as follows: initial denaturation step at 98°C for 1 minute, followed by 25 cycles of: denaturation step at 98°C for 10 seconds, annealing step at 50°C for 30 seconds and elongation step at 72°C for 30 seconds; finishing with the extension step at 72°C for 4 minutes. For fungal analyses, the ITS1 fungal region was amplified using the primers ITS1f and ITS2 augmented with multiplexing barcodes (Smith & Peay 2014). Each reaction mixture contained 1 µl of sample, 1 μl of forward primer (5 μM) and 12.5 μl Phusion High-Fidelity PCR Master Mix. PCR conditions were as follows: initial denaturation step at 98°C for 30 s, followed by 30 cycles of: 98°C for 30 s, 55°C for 30 s, 72°C for 30 s; and an additional extension step of 72°C for 10 min.

Samples that failed to produce PCR product were subject to repeated soil extraction and PCR. However, 7 samples for bacteria (1 from W_0_ and W_med_, 2 from W_high_ and W_extr_ in LWG; 1 from W_high_ in SWG) and 6 samples for fungi (1 from W_high_, 2 from W_extr_, in LWG, 3 from W_high_ in SWG) still failed to produce usable PCR products and these were excluded from further analyses. Successful amplification products were purified using the AmpureXP method (Beckman Coulter, Brea, CA, United States) and normalized to equimolar concetrations before pooling into a single library, for fungi and bacteria separately. Gel extraction of the pooled library was performed for size selection and additional purification using QIAquick Gel Extraction Kit (Qiagen, Venlo, the Netherlands). Libraries were quantified with real-time PCR (KAPA Library Quantification Kits, Kapa Biosystems, Wilmington, MA, USA).

The libraries were sequenced on the Illumina MiSeq platform (Illumina Inc; San Diego, CA, USA) with 150 cycles, for forward and reverse reads for bacteria and 300 cycles in the forward direction for fungi. The reproducibility of sample preparation and sequencing was tested by sequencing a small number of technical replicates (DNA isolated from the same samples but subjected to independent PCR reactions with distinct indices) (Fig. S1).

### 2.3 Quality filtering and bioinformatics analysis

The first part of bioinformatics analysis on bacterial sequences was performed using the USEARCH software (Edgar 2013). Paired-end reads were merged and primer sequences removed. The sequences were subsequently quality filtered (maximum expected error 0.05) leaving a total of 6.4M high-quality sequences. Following dereplication and singleton removal a set of OTU representative sequences (97% similarity) was constructed using the UPARSE-OTU algorithm (Edgar, 2013). After chimera removal (leaving 19,423 non-chimeric OTUs) all original reads were mapped to the non-chimeric OTUs using the USEARCH algorithm with global alignments with the identity threshold of 0.97, yielding an OTU table. From this point on, all subsequent steps were performed with QIIME software (Caporaso et al. 2010b). OTUs were aligned to the Green Genes February 2011 database (DeSantis et al. 2006) using the PyNAST algorithm (Caporaso et al. 2010a). To avoid library size-related artefacts, a subsampled OTU table was created by random sampling of the original OTU table. In this step, samples that contained fewer sequences than the requested depth (7000) were omitted from the output OTU tables. This rarefaction depth included all but two samples (from W_0_ and W_high_ group in LWG) which had a significantly lower amount of sequences than other samples, and whose reliability was therefore questionable. Taxonomic identity of each OTU was identified based on the 97% Green Genes database (release 13_8) using the RDP classifier (Wang et al. 2007).

Fungal sequences were analysed using the USEARCH software following the UPARSE pipeline (Edgar, 2013). After trimming to 200 bp, the sequences were quality filtered (maximum expected error 0.01), leaving a total of 4.15 M sequences. The reverse primers were removed and N’s were added up to 200 bp for efficient clustering of OTUs. Singleton sequences were removed and all others were clustered to 97% similarity. Chimeras were filtered *de novo* as well as through the UNITE database of ITS1 sequences as implemented in UCHIME, resulting in a total of 3,618 non-chimeric OTUs. Representative sequences for each OTU were aligned to all fungal representative species in the UNITE database (Kõljalg et al. 2005)(release date 01.08.2015), using the BLAST algorithm with default settings. The resulting hits were assigned according to taxa selecting hits with the lowest E-value and a minimum alignment length of 75 bp. OTUs were classified to taxonomic levels depending on identity percentages with UNITE taxa using the following thresholds: > 90% identity level corresponds to genus, > 85% to family, as in Tedersoo et al. (2014). OTUs were subsequently assigned to functional groups if the genus was successfully matched with one of the genera with known lifestyles in Tedersoo et al. (2014). In the cases when genus level was unknown, lifestyle was assigned at family level if more than 80% of genera within that family (represented by more than 3 genera) belonged to the same lifestyle. All original sequences were mapped against these OTUs with a similarity threshold of 97% and assembled in an OTU table. The number of reads per sample was then rarefied to the minimum number of reads of 5,274.

### 2.4 Statistical analyses

Nonmetric multidimensional scaling (NMDS) was performed to visualize the overall differences in microbial community composition. Generalized additive models were used to evaluate the correlation between the ordination of samples based on two-dimensional NMDS and the temperature elevation as a continuous variable ranging from +0°C to +19°C. The differences between microbial communities from the soil exposed to different warming levels (W_low_, W_med_, W_high_/W_high_’, W_extr_) and the unwarmed soil (W_0_), were quantitatively evaluated through pair-wise PERMANOVA analysis (Anderson 2001). Bonferroni correction was applied to each separate test to adjust P values for multiple testing in pair-wise PERMANOVA. All multivariate analyses were based on Bray-Curtis (BC) distances to allow comparability between bacterial and fungal datasets, but the results were robust to the choice of distance metric (data not shown). The correlation between temperature elevations and the BC distances of fungal/bacterial communities from different plots was quantified using a Mantel test. For all statistical tests, OTU abundances were log transformed.

The change in the relative abundance of dominant bacterial high-level taxa / fungal functional groups (with the amount of sequences greater than 2% of the total number) at different warming levels was tested using ANOVA (followed by posthoc Tukey tests in the cases of significant results). When necessary, the data were transformed (using log or Box-Cox transformations) to meet the normality assumption. P values were in all cases adjusted for multiple testing using Bonferroni correction.

All statistical analyses were conducted using R statistical software (R Core Team 2015) using base packages and vegan (Oksanen et al. 2015).

## 3. RESULTS

### 3.1 Overall community composition along warming gradients

The results of PERMANOVA analysis showed that, in the LWG, bacterial community composition was significantly affected only by the highest temperature elevations W_extr_ (+15°C to +19°C). Temperature elevations below this range (i.e. up to +9 °C) did not lead to bacterial community compositions differing from ambient soils (Table 1, Fig 2a, Fig. 2c). It should be noted, however, that group W_high_ (+6°C to +9°C) only had two observations and the comparison with this group has therefore a higher uncertainty (Table 1). In SWG, bacterial communities from W_high_’ (+7°C to +11°C) and W_extr_ (+15°C to +19°C) warming levels differed significantly from the ambient group (R^2^=10%, P<0.01and R^2^=17%, P<0.01, respectively), while the communities at lower temperature elevations were not significantly different to those from ambient soils (Table 1, Fig. 2b, Fig. 2c).

**Table 1.**
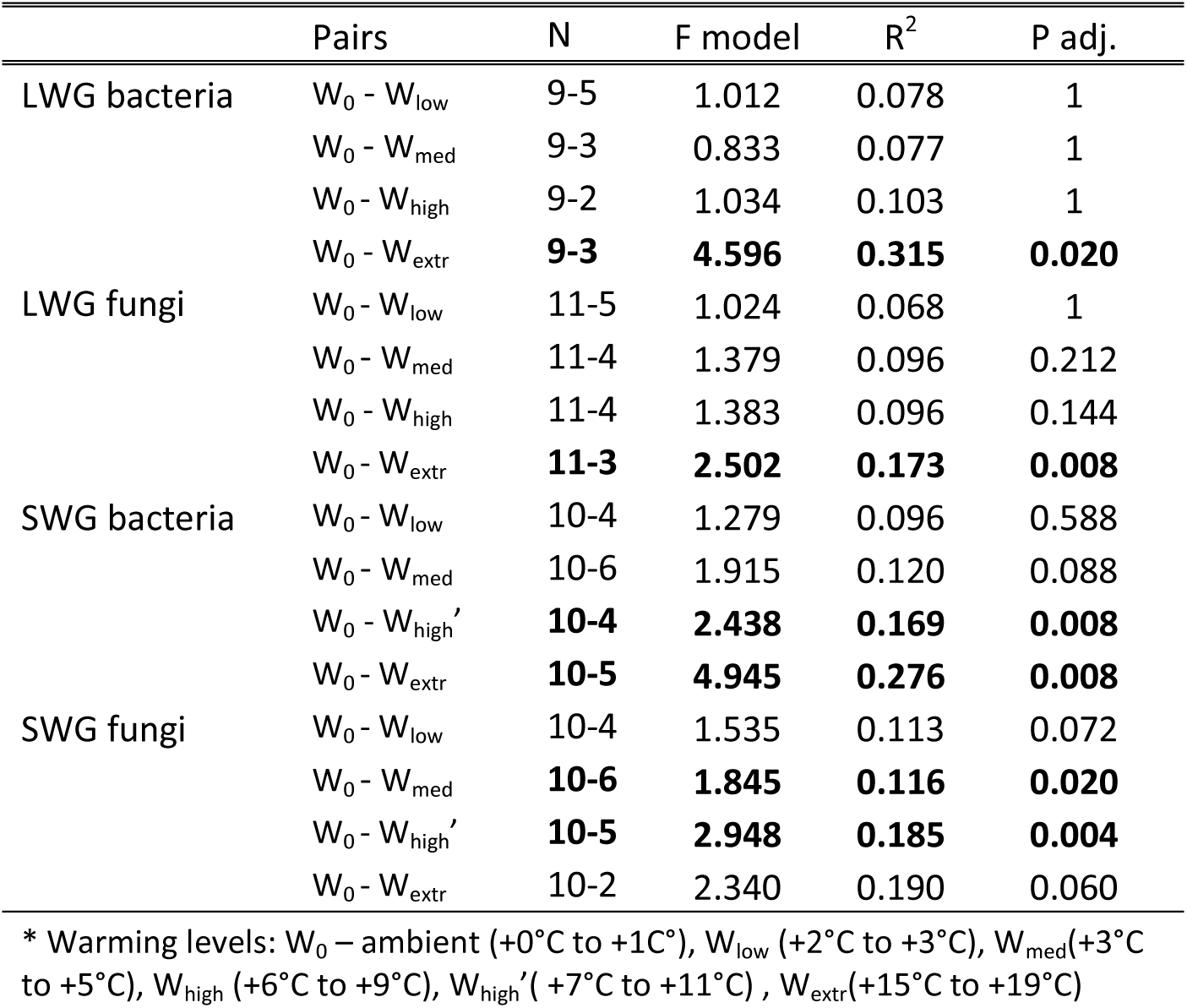
The results of pairwise PERMANOVA analyses between the communities from ambient soil temperatures and the communities from increased soil temperatures, in the long-term warmed (LWG) and the short-term warmed (SWG) grassland.

**Figure 2.**
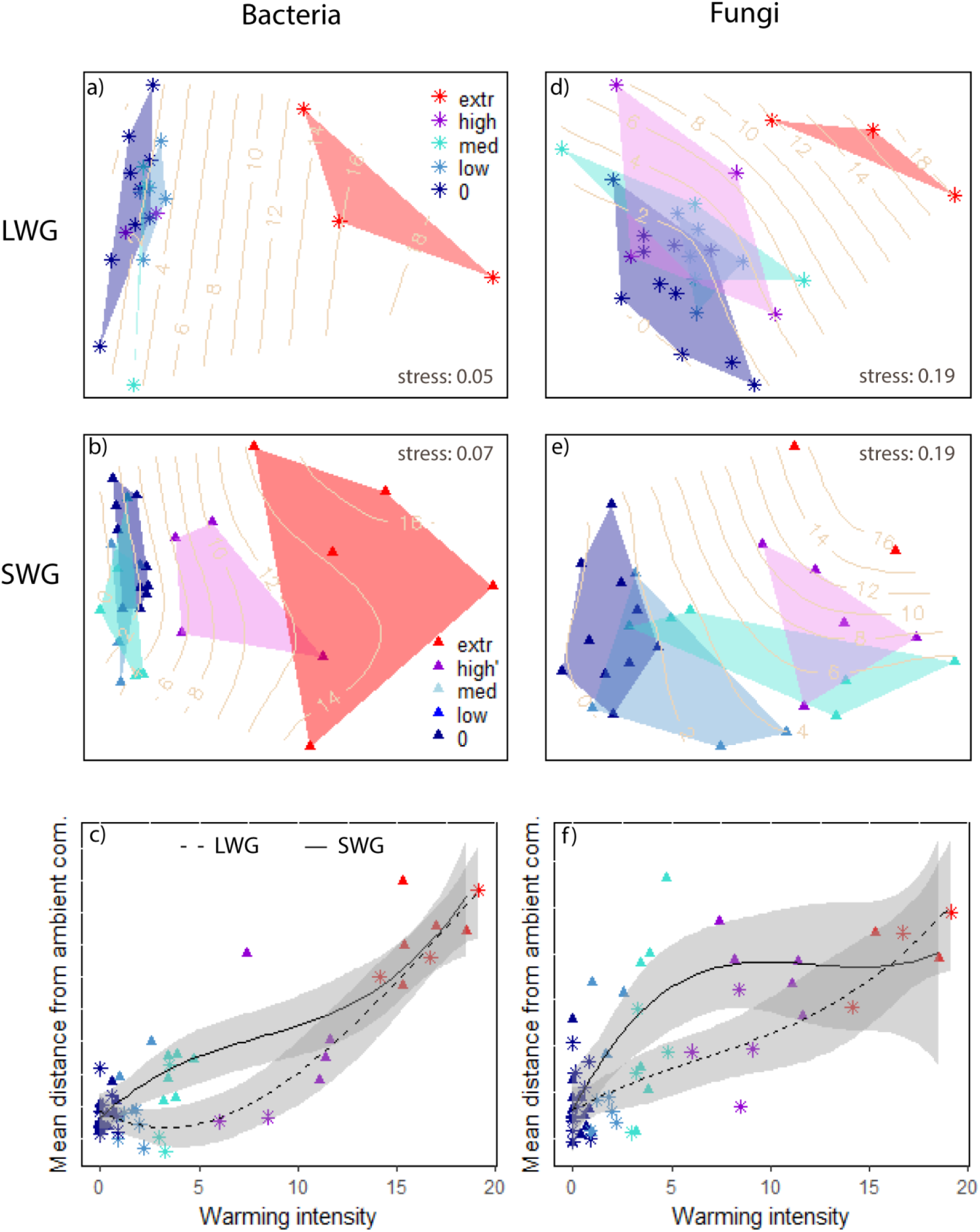
**Top**: NMDS ordination plots for: bacterial community composition in a) the long-term warmed grassland(LWG) and b) the short-term warmed grassland (SWG); and fungal community composition in d) LWG and f) SWG. Points (samples) and the corresponding polygons are coloured according to warming levels W [0 – ambient (+0°C to +1C°), low (+2°C to +3°C), med (+3°C to +5°C), high (+6°C to +9°C), high’ (+7°C to +11°C), extr (+15°C to +19°C)]. Isolines represent fitted smooth surface of continuous different warming intensities. **Bottom:** Mean distances (based on Bray-Curtis dissimilarity between the communities from each sample and the ambient communities along the temperature gradient for c) bacterial comunities and f) fungal communities in LWG and SWG. The lines represent smoothing splines (3^rd^ order polynomial model), dashed line = LWG, full line = SWG. Shaded area represents 95% confidence intervals of the fitted lines calculated using the predict method.

For fungal communities in the LWG, the significant change was also found only for the W_extr_ (R^2^=17%, P<0.01) (Table 1, Fig 2d, Fig. 2f). In SWG, fungal communities were significantly affected by warming levels W_med_ (+3°C to +5°C) and W_high_’(+7°C to +11°C) (R^2^=12%, P=0.02 and R^2^=19%,P=0.004; respectively), while the difference for W_low_ (+1°C to +3°C) and W_extr_ (+15°C to +19°C) was borderline significant (R^2^=11%, P=0.07 and R^2^=19%, P=0.06; respectively) (Table 1, Fig. 2f, Fig. 2d). Absence of a significant effect in the latter likely results from low statistical power due to small sample size (n=2), especially when considered that the effect size for this group is relatively high and that they are clearly separated from the ambient communities based on NMDS ordination (Fig. 2e).

Based on a generalized additive model, there was a significant relationship (P<0.001) between the ordination axes in NMDS and temperature elevations: in LWG (R^2^=86% and R^2^=88%, for bacteria and fungi, respectively) and in SWG (R^2^=84%, R^2^=83%, for bacteria and fungi, respectively). This indicates that the ordination patterns observed from the plots (Figure) can to a large extent be explained by the differences in soil temperatures. In addition, a Mantel test indicated that community dissimilarities between the samples were significantly related to the differences in soil temperatures both in LWG (R^2^=70%, P<0.01 and R^2^=54%, P<0.01) and in SWG (R^2^=60%, P=0.001 and R^2^=42%, P=0.001, for bacteria and fungi, respectively).

### 3.2 Microbial taxa/functional-groups along warming gradients

The most dominant bacterial phyla in both LWG and SWG were: Proteobacteria (Alphaproteobacteria 13%/15% of the reads; Beta-proteobacteria 8%/5%; Deltaproteobacteria 6%/6%; Gammaproteobacteria 2%/2%), Acidobacteria (25%/18%), Actionobacteria (11%/21%) and Chloroflexi (6%/6%) in LWG and SWG, respectively. Other phyla with more than 2% of total number of sequences were: Verrucomicrobia, Gemmatimonadetes, Bacteroidetes, Firmicutes and Nitrospirae. The list of all bacterial phyla in LWG and SWG can be found in Table S1.

Filamentous saprotrophs were by far the most dominant fungal functional group (66% and 62% in LWG and SWG, respectively), followed by arbuscular mycorrhizal (AM) fungi (6% and 5%) and yeasts (2% and 7%). Other functional groups: ectomycorrhizal fungi, white rot saprotrophs, plant pathogens and mycoparasites accounted for less than 2% of total number of sequences (Table S2).

Based on ANOVA analysis, there was one dominant bacterial high-level taxon (comprising more than 2% of all sequences) that differed significantly among different warming levels in LWG (Betaproteobacteria) and there were three in SWG (Betaprotobacteria, Bacteroidetes, Chloroflexi)(Table S3). A post hoc test (Table S4) showed that in LWG, the relative abundance of Betaproteobacteria significantly decreased only at W_extr_ (P=0.01) compared to the ambient (Fig. 3). In SWG the relative abundance of Betaproteobacteria and Bacteroidetes was significantly decreased at W_high_’ and W_extr_ (P<0.001), while the relative abundance of Chloroflexi was significantly increased at W_extr_ (P<0.001) compared to the ambient.

For fungal communities, a significant increase in the relative abundance of AM fungi compared to the ambient was found for the warming level W_extr_ (P<0.001) in LWG and for warming levels W_high_’ and W_extr_ (P<0.05) in SWG. The relative abundance of filamentous saprophytes decreased significantly at W_extr_ (P<0.001) in LWG and at W_high_’ and W_extr_ (P=0.01 and P<0.001, respectively) in SWG (Fig. 3). The third dominant group (yeasts) did not show a prominent pattern along the warming gradient (Table S3).

**Figure 3.**
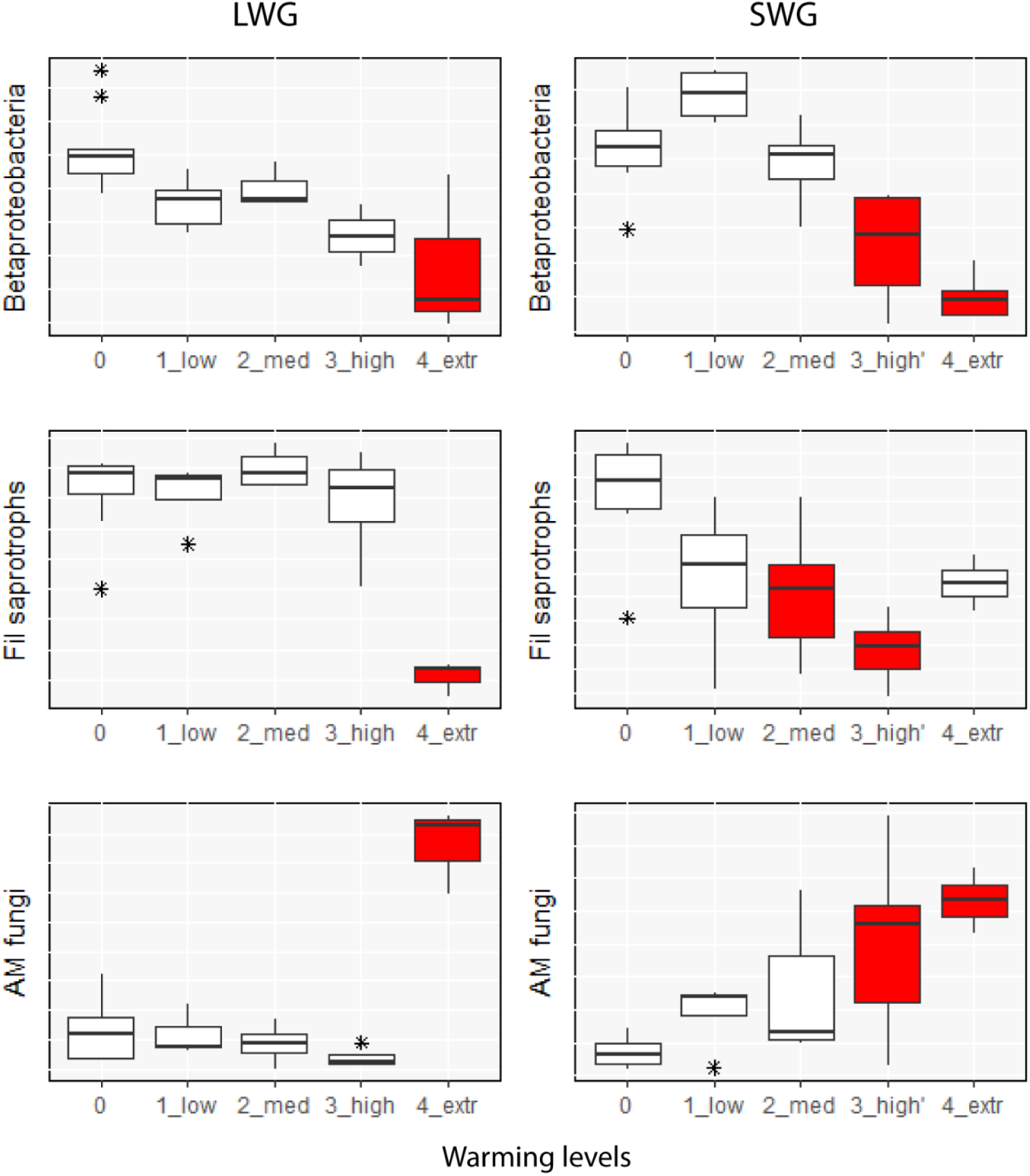
The change in the relative abundance (% of the total amount of sequences in a sample) of Betaproteobacteria and fungal functional groups (AM fungi and filamentous saprotrophs) in the long-term warmed grassland (LWG; left) and the short-term warmed grassland (SWG; right). Only the microbial groups that differed significantly between different warming levels in both grasslands are shown. Warming levels: W [0 – ambient (+0°C to +1C°), 1_low (+2°C to +3°C), 2_med (+3°C to +5°C), 3_high (+6°C to +9°C), 3_high’ (+7°C to +11°C), 4_extr (+15°C to +19°C)]. Red box plots indicate warming levels where the relative abundance of a given microbial group was significantly different from the relative abundance at ambient temperature based on post-hoc Tukey test.

## 4. DISCUSSION

Substantial efforts have been invested into research of the warming effects on microbial community composition and functioning (Griffiths & Philippot 2013). However, there have been very few studies investigating the effect of the exposure period on microbial responses to warming, despite evidence that the timescale of warming could be an important factor in detecting significant responses (Rinnan et al. 2007; DeAngelis et al. 2015). The present study took advantage of natural geothermal gradients of different ages (5-7 years and >50 years) and high-resolution DNA sequencing technique to test the hypothesis that prolonged exposure intensifies the effect of warming on soil microbial community composition so that it can be detected at lower temperature elevations.

### 4.1 Long-term versus short-term effects of warming

Community profiles from amplicon sequencing of bacterial and fungal rRNA genes showed that significant changes in community composition relative to ambient were only observed above +9°C in the long-term and at +7°C to +11°C / +3°C to +5°C in the short-term for bacteria and fungi, respectively (Fig. 2). Collectively, these results refute our hypothesis that long-term exposure intensifies microbial community responses to warming, since there was no evidence that detectable changes from ambient communities occurred at lower temperature elevations in the long-term warmed grassland than in the short-term warmed grassland. It is therefore unlikely that the responses observed in short-term warming experiments (5-7 years) in high-latitude grasslands would underestimate the long-term effect of soil warming on microbial community composition. On the other hand, against our expectations, there was some evidence that changes in microbial community composition occurred at lower temperature elevations in the short-term compared to the long-term warmed grassland (Fig. 1c and Fig. 1f). However, this cannot be explicitly claimed for bacteria due to the difference in warming range in the W_high_ group, which was slightly lower in LWG (+6°C to +9°C) than in SWG (+7°C to +11°C), and the fact that there was a substantial gap in data within this warming range. This pattern was, however, more strongly supported for fungal communities since the differences in responses between the two grasslands were apparent at lower warming ranges (between +3°C and +5°C) where the larger amount of data produces higher certainty in the result (Fig. 1f).

We propose the following potential explanations for the observed patterns: (i) after an initial shift in response to warming, microbial communities reverted towards their pre-disturbance composition over the years, except at very high temperatures, where a new (stable) state was established; (ii) the inconsistencies between warming effects in the two grasslands are a consequence of site-specific effects rather than the differences in exposure period.

The proposition that microbial community shifts could be transient after a long period of warming contradicts a number of studies which reported that the effects of warming were detected even after many years (up to 20) of continuous exposure (Allison & Martiny 2008; Frey et al. 2008; Deslippe et al. 2012; Luo et al. 2014; DeAngelis et al. 2015). However, none of these studies examined a warming period as long as the one found in LWG (more than 50 years, possibly several centuries). It is possible that the warming had initially altered some components of the soil ecosystem (e.g. the availability of nutrients, abundance of predators, root density or plant composition), which then led to a substantial change in soil microbial community composition. However, if shifts in these intermediate drivers were transient over longer time scales (e.g. predator or plant communities re-established due to the colonization of warmed areas by more tolerant species with similar functions), microbial community composition could potentially have changed back towards the original state accordingly. In line with this, at the ForHot site, it has been observed that collembolan communities were affected more strongly by short-term than long-term warming (Holmstrup et al. subm.). Similar results were found for nematode and plant communities (Ilieva-Makulec, Leblans; pers. comm.). This would imply that warming itself is not the single most important factor structuring microbial communities, but some other warming-induced changes in biotic and/or abiotic environmental factors might play a substantial role. Indeed, a growing body of evidence supports the assumption that certain factors which are typically influenced by warming, such as: nutrient availability (Koyama et al. 2014) plant productivity or diversity (Zak et al. 2003; Zhang et al. 2005) or other food web members such as nematodes (Ruess et al. 1999) oribatid mites (Maraun et al. 1998) and collembolan (Tiunov & Scheu 2005), have an important impact on soil microbial communities.

When considering the apparent differences in microbial community responses between the long-term and the short-term warmed grassland, it should be emphasized that our study did not follow one single system through time, but two different systems with different exposure periods to warming. It is therefore possible that the observed differences in the community temperature responses between LWG and SWG could be attributed to the differences in particular environmental conditions in these two grasslands. The grasslands are located within the distance of 3 km, they are subject to the same climatic conditions, dominated by the same plant species and they have the same type of soil with similar pH conditions (Sigurdsson et al. 2016). Nevertheless, microbial communities at ambient temperatures in LWG differed significantly from the corresponding communities in SWG (Fig. S2). If these different microbial communities (or their site-specific drivers) respond differently to the same temperature elevation, this would preclude attributing such an effect to warming period per se.

### 4.2 The response of high-level microbial groups to warming

In general, it was observed that the responses of individual bacterial OTUs were not consistent within higher taxonomic classifications since few high-level taxonomic groups showed a significant trend along the temperature gradient (Table S3). Therefore, the changes in bacterial community composition with warming were probably a result of shifts at lower taxonomic levels. However, the decrease in relative abundance of members of the Betaproteobacteria subphylum corresponded to the pattern in overall bacterial community composition along the gradient in both grasslands (Fig. 2c, Fig. 3). This implies that the observed patterns in bacterial community composition may, to a certain extent, be explained by the decrease in abundance of Betaproteobacteria. Several other studies have reported a consistent response to environmental drivers, including temperature elevation, for Betaproteobacteria (Fierer et al. 2007; Rui et al. 2015; Zhang et al. 2016) indicating that members of this group may largely share environmental requirements.

For soil fungi the significant changes in abundance of filamentous saprotrophs and AM fungi roughly coincide with the changes in overall fungal community composition along the two gradients (Fig. 2f, Fig. 3). Since filamentous saprotrophs and AM fungi together represent the majority of the total amount of fungal sequences found in the soil samples (72% and 67% for LWT and SWT, respectively), it can be concluded the changes in overall fungal community composition are largely the consequence of changes in these two groups along the gradient. These results suggest that warming may induce shifts from free living saprotrophs towards AM fungi. It has been proposed that the shifts in fungal functional groups may alter some ecosystem functions such as soil carbons storage (Treseder & Lennon 2015). In line with our results, Rudgers et al. (2014) have reported a tendency of AM fungi to increase, and non-AM filamentous fungi to decrease after 20 years of experimental warming by 2°C. In contrast, Heinemeyer et al. (2004), reported no consistent warming effects on AM fungi after one season of exposure to approximately +3°C.

### 4.3 Sensitivity of microbial communities to climate warming

Our results provide insights into the sensitivity of soil microbial communities under predicted climate warming in high-latitude systems (+2.2°C to +8.3°C, by the end of century (IPCC 2013)) while taking into account different exposure periods and warming intensities. We argue that low-intensity warming, at levels typically applied in static warming experiments (between +1°C and +5°C), is not likely to significantly alter bacterial and fungal community composition in this subarctic grassland system over the long term. In the short-term, fungi but not bacteria were already affected by warming intensities of +3°C, suggesting that future temperature increases may transiently alter fungal community composition. This differential response between fungi and bacteria could be interpreted as reflecting differing temperature sensitivities and/or recovery times of the respective communities. However, we caution against over-interpreting this pattern as it could be a consequence of the two year gap in sampling within the current study.

The lack of consistent conclusions regarding the sensitivity of soil microbial communities to warming in different studies, could arise both from ecosystem dependency (Allison & Martiny 2008; Pold & DeAngelis 2013; Cregger et al. 2014; Zhang et al. 2016) and inconsistencies in methodological approaches. This study is unique in that it focuses on the effect of warming on both fungal and bacterial communities along temperature gradients, unlike most other studies that use single-temperature manipulation treatments and are thus not able to evaluate the effect of different warming magnitudes (Thompson et al. 2013). Given that high-latitude systems will potentially be subject to a higher intensity of warming than most other regions (Serreze et al. 2000; MacDonald 2010, IPCC 2013), it is particularly important to get a better mechanistic understanding of the effect of different temperature elevations in these ecosystems. On the other hand, it must be emphasised that geothermal soil warming, itself, is not entirely comparable to the warming resulting from climate change, particularly because the aboveground systems do not experience substantial temperature increase (Sigurdsson et al. 2016). This is, however, a common issue for most studies investigating the effects of warming on soil systems. In terms of differences arising from community profiling techniques, we utilized a high resolution, next generation sequencing of molecular markers that allows detection of changes in low taxonomic levels, unlike commonly used biochemical approaches such as PLFA analysis.

## CONCLUSION

The present study demonstrated that, contrary to expectations, long-term exposure did not intensify the effect of warming on microbial community composition. We thus conclude that the effects of warming on microbial community composition observed in short-term experiments (with the duration of 5-7 years,) in subarctic grasslands are not likely to underestimate effects over larger time scales – in contrast, our results suggest a risk of overestimation. The significant influence of warming was observed at medium (+3°C to +5°C) and upper (> +7°C) ranges of predicted temperature increase in next 100 years (+2.2°C to +8.3°C), for fungi and bacteria, respectively. Given that these effects may not persist in the long-term, microbial communities in high-latitude ecosystems could be less sensitive to climate warming than previously expected. This study highlights the value of natural warming gradients as study systems which can provide valuable information on the effect of different warming intensities on microbial communities, an important topic that has received little attention thus far.

## Author contributions

BDS and NIWL designed the experiment. NIWL and DR performed the field work. JTW, EV and DR performed the lab work, bioinformatic and statistical analyses. JTW, EV, SV and DR interpreted the data. DR, JTW and EV wrote the first draft of the manuscript and all co-authors contributed to the final version.

## ACKNOWLEDGEMENTS

The authors acknowledge E. Oerlemans and S. Lebeer for generously providing molecular facilities, A. Meire and S. Dauwe for assistance with field sampling and J. de Gruyter for DNA extractions. Sequencing was performed at the Center for Medical Genetics of the University of Antwerp. JTW and SV were supported by a postdoctoral fellowship grant of the Research Foundation–Flanders (FWO) and NIWL was supported by a FWO aspirant grant. BDS was supported by Icelandic Research Council. We also acknowledge support of ClimMani COST Action (ES1308) from the European Research Council grant ERC-SyG-610028 IMBALANCE-P.

## SUPPORTING INFORMATION

**Table S1** The percentage of sequences belonging to different bacterial phyla

**Table S2** The percentage of sequences belonging to different fungal functional groups

**Table S2** The results of ANOVA analyses

**Table S3** The results of Tukey test

**Figure S1** NMDS ordination of technical replicates

**Figure S2** NMDS ordination showing the differences between ambient communities

## References

Allison SD & Martiny, JBH (2008). Resistance, resilience, and redundancy in microbial communities. Proc Nat Acad Sci USA, 105, 11512–11519.

Allison SD, McGuire KL & Treseder KK (2010). Resistance of microbial and soil properties to warming treatment seven years after boreal fire. Soil Biol Biochem, 42, 1872–1878.

Anderson MJ (2001). A new method for non-parametric multivariate analysis of variance. Austral Ecology, 26, 32–46.

Bartram AK, Lynch MD, Stearns JC, Moreno-Hagelsieb G & Neufeld JD (2011). Generation of multimillion-sequence 16S rRNA gene libraries from complex microbial communities by assembling paired-end Illumina reads. Appl Environ Microb, 77, 3846–3852.

Blankinship JC, Niklaus PA & Hungate, BA (2011). A meta-analysis of responses of soil biota to global change. Oecologia, 165, 553–565.

Caporaso GJ, Bittinger K, Bushman FD, DeSantis TZ, Andersen GL, & Knight R (2010a). PyNAST: a flexible tool for aligning sequences to a template alignment. Bioinformatics, 26, 266–267.

Caporaso GJ, Kuczynski J, Stombaugh J, Bittinger K, Bushman FD, Costello, EK, et al. (2010b). QIIME allows analysis of high-throughput community sequencing data. Nat Methods, 7, 335–336.

Contosta, A.R., Frey, S.D. & Cooper, A.B., 2015. Soil microbial communities vary as much over time as with chronic warming and nitrogen additions. Soil Biol Biochem, 88, 19–24.

Cregger MA, Sanders NJ, Dunn RR & Classen AT (2014). Microbial communities respond to experimental warming, but site matters. PeerJ, 2, p.e358.

De Boeck HJ, Vicca S, Roy J, Nijs I, Milcu A, Kreyling Jet al. (2015). Global change experiments: challenges and opportunities. BioScience, 65, 922–931.

DeAngelis, KM, Pold G, Begüm D, van Diepen LTS, Varney, RM, Blanchard JLet al. (2015). Long-term forest soil warming alters microbial communities in temperate forest soils. Front Microbiol, 6, 104.

DeSantis TZ, Hugenholtz P, Larsen N, Rojas M, Brodie EL, Keller K, et al. (2006). Greengenes, a chimera-checked 16S rRNA gene database and workbench compatible with ARB. Appl Environ Microb, 72, 5069–72.

Deslippe JR, Hartmann M, Simard SW & Mohn WW 2012. Long-term warming alters the composition of Arctic soil microbial communities. FEMS Microbiol Ecol, 82, 303–315.

Edgar, RC (2013). UPARSE: highly accurate OTU sequences from microbial amplicon reads. Nat Methods, 10, 996–998.

Fierer N, Bradford MA. & Jackson RB (2007). Toward an ecological classification of soil bacteria. Ecology, 88, 1354–1364.

Frey SD, Drijber R, Smith H & Melillo J (2008). Microbial biomass, functional capacity, and community structure after 12 years of soil warming. Soil Biol Biochem. 40, 2904–2907.

Griffiths BS & Philippot L (2013). Insights into the resistance and resilience of the soil microbial community. FEMS Microbiol Rev, 37, 112–129.

Heinemeyer A, Ridgway KP, Edwards EJ, Benham DG, Young JP & Fitter AH (2004). Impact of soil warming and shading on colonization and community structure of arbuscular mycorrhizal fungi in roots of a native grassland community. Glob Change Biol, 10, 52–64.

IPCC, 2013: Climate Change 2013: The Physical Science Basis. Contribution of Working Group I to the Fifth Assessment Report of the Intergovernmental Panel on Climate Change [Stocker TF, Qin D, Plattner G-K, Tignor M, Allen SK, Boschung J, Nauels, A, Xia Y, Bex V & Midgley PM (eds.)]. Cambridge University Press, Cambridge, United Kingdom and New York, NY, USA, 1535 pp.

Karhu K, Auffret MD, Dungait JA, Hopkins DW, Prosser JI, Singh BKet al. (2014). Temperature sensitivity of soil respiration rates enhanced by microbial community response. Nature, 513, 81–84.

Kõljalg U, Larsson KH, Abarenkov K, Nilsson RH, Alexander IJ, Eberhardt Uet al. (2005). UNITE: a database providing web-based methods for the molecular identification of ectomycorrhizal fungi. New Phytologist, 166, 1063–1068.

Koyama A, Wallenstein MD, Simpson RT & Moore JC (2014). Soil bacterial community composition altered by increased nutrient availability in Arctic tundra soils. Front Microbiol, 5, 516.

Luo CW, Rodriguez-R LM, Johnston ER, Wu LY, Cheng L, Xue Ket al. (2014). Soil microbial community responses to a decade of warming as revealed by comparative metagenomics. Appl Environ Microb, 80, 1777–1786.

MacDonald, GM (2010). Global warming and the Arctic: a new world beyond the reach of the Grinnellian niche? Journal of Experimental Biology, 213, 855–861.

Männistö MK, Tiirola M & Häggblom MM (2007). Bacterial communities in Arctic fjelds of Finnish Lapland are stable but highly pH-dependent. FEMS Microbiol Ecol, 59, 452–65.

Maraun M, Visser S & Scheu S (1998). Oribatid mites enhance the recovery of the microbial community after a strong disturbance. Applied Soil Ecology, 9, 175–181.

O'Gorman EJ, Benstead JP, Cross WF, Friberg N, Hood JM, Johnson PWet al. (2014). Climate change and geothermal ecosystems: Natural laboratories, sentinel systems, and future refugia. Glob Change Biol, 20, 3291–3299.

Oksanen J, Blanchet FG, Friendly M, Kindt R, Legendre P, McGlinn Det al. (2015). Vegan: community ecology package. R package version 2.4–0.

Penton CR, StLouis D, Cole JR, Luo Y, Wu L, Schuur EGet al. (2013). Fungal diversity in permafrost and tallgrass prairie soils under experimental warming conditions. Appl Environ Microb, 79, 7063–7072.

Pold G & DeAngelis, KM (2013). Up against the wall: The effects of climate warming on soil microbial diversity and the potential for feedbacks to the carbon cycle. Diversity, 5, 409–425.

R Core Team, 2015. R: A language and environment for statistical computing. R Foundation for Statistical Computing.

Rinnan R, Michelsen A, Bååth E & Jonasson S (2007). Fifteen years of climate change manipulations alter soil microbial communities in a subarctic heath ecosystem. Glob Change Biol, 13, 28–39.

Rousk J, Smith AR & Jones, DL (2013). Investigating the long-term legacy of drought and warming on the soil microbial community across five European shrubland ecosystems. Glob Change Biol, 19, 3872–3884.

Rudgers JA, Kivlin SN, Whitney KD, Price MV, Waser NM & Harte J (2014). Responses of high-altitude graminoids and soil fungi to 20 years of experimental warming. Ecology, 95, 1918–1928.

Ruess L, Michelsen A, Schmidt IK & Jonasson S (1999). Simulated climate change affecting microorganisms, nematode density and biodiversity in subarctic soils. Plant and Soil, 212, 63–73.

Rui J, Li J, Wang S, An J, Liu WT, Lin Q, Yang Yet al. (2015). Responses of bacterial communities to simulated climate changes in alpine meadow soil of the Qinghai-Tibet Plateau. Appl Environ Microb, 81, 6070–6077.

Rustad L (2001). Global change: Matter of time on the prairie. Nature, 413, 578–579.

Schindlbacher A, Rodler A, Kuffner M, Kitzler B, Sessitsch A & Zechmeister-Boltenstern S (2011). Experimental warming effects on the microbial community of a temperate mountain forest soil. Soil Biol Biochem, 43, 1417–1425.

Serreze MC, Walsh JE, Chapin FS, Osterkamp T, Dyurgerov M, Romanovsky V, et al. (2000). Observational evidence of recent change in the northern high-latitude environment. Climatic Change, 46, 159–207.

Sigurdsson BD, Leblans NI, Dauwe S, Guðmundsdóttir E, Gundersen P, Gunnarsdóttir GE, et al. (2016). Geothermal ecosystems as natural climate change experiments: The ForHot research site in Iceland as a case study. Icel.agric.sci, 29, 53–71.

Smith DP & Peay KG (2014). Sequence depth, not PCR replication, improves ecological inference from next generation DNA sequencing. PLoS ONE, 9, 90234.

Strickland MS, Lauber C, Fierer N & Bradford MA (2009). Testing the functional significance of microbial community composition. Ecology, 90(2), 441–451.

Tedersoo L, Bahram M, Põlme S, Kõljalg U, Yorou NS, Wijesundera R, et al. (2014). Global diversity and geography of soil fungi. Science, 346, 1256688.

Thompson RM, Beardall J, Beringer J, Grace M & Sardina P (2013). Means and extremes: building variability into community-level climate change experiments. Ecol Letts, 16, 799–806.

Tiunov, AV & Scheu S (2005). Arbuscular mycorrhiza and Collembola interact in affecting community composition of saprotrophic microfungi. Oecologia, 142, 636–642.

Treseder KK. & Lennon JT (2015). Fungal traits that drive ecosystem dynamics on land. Microbiol Mol Biol Rev, 79, 243–262.

Wang Q, Garrity GM, Tiedje JM & Cole JR (2007). Naive bayesian classifier for rapid assignment of rRNA sequences into the new bacterial taxonomy. Appl Environ Microb, 73, 5261–5267.

Weedon JT, Kowalchuk GA, Aerts R, van Hal JR, van Logtestijn RKSP, Taş Net al. (2012). Summer warming accelerates sub-arctic peatland nitrogen cycling without changing enzyme pools or microbial community structure. Glob Change Biol, 18, 138–150.

Woodward G, Dybkjaer JB, Ólafsson JS, Gíslason GM, Hannesdóttir ER & Friberg N (2010). Sentinel systems on the razor’s edge: effects of warming on Arctic geothermal stream ecosystems. Glob Change Biol, 16, 1979–1991.

Xiong J, Sun H, Peng F, Zhang H, Xue X, Gibbons SMet al. (2014). Characterizing changes in soil bacterial community structure in response to short-term warming. FEMS Microbiol Ecol, 89, 281–292.

Xu G, Chen J, Berninger F, Pumpanen J, Bai J, Yu Let al. (2015). Labile, recalcitrant, microbial carbon and nitrogen and the microbial community composition at two Abies faxoniana forest elevations under elevated temperatures. Soil Biol Biochem, 91, 1–13.

Zak DR, Holmes WE, White DC, Peacock AD & Tilman D (2003). Plant diversity, soil microbial communities, and ecosystem function: Are there any links? Ecology, 84, 2042–2050.

Zhang K, Shi Y, Jing X, He JS, Sun R, Yang Yet al. (2016). Effects of short-term warming and altered precipitation on soil microbial communities in alpine grassland of the Tibetan Plateau. Front microbiol, 7, 1032.

Zhang W, Parker KM, Luo Y, Wan S, Wallace LL & Hu S (2005). Soil microbial responses to experimental warming and clipping in a tallgrass prairie. Glob Change Biol, 11, 266–277.

Zogg GP, Zak DR, Ringelberg DB, White DC, MacDonald NW & Pregitzer KS (1997). Compositional and functional shifts in microbial communities due to soil warming. Soil Sci Soc Am J, 61, 475–481.

Þorbjörnsson D, Sæmundsson K, Kristinsson S & Kristjánsson B (2009). Suðurlandsskjálftar 29. maí 2008: Áhrif á grunnvatnsborð, hveravirkni og sprungumyndun (The south Iceland earthquake of May 29, 2008: Effects on ground water table, geothermal activity, and fractures). Report Iceland GoeSurvey, iSOR-2009/028, 42.

